# FRETpredict: A Python package for FRET efficiency predictions using rotamer libraries

**DOI:** 10.1101/2023.01.27.525885

**Authors:** Daniele Montepietra, Giulio Tesei, João M. Martins, Micha B. A. Kunze, Robert B. Best, Kresten Lindorff-Larsen

## Abstract

Here, we introduce FRETpredict, a Python software program to predict FRET efficiencies from ensembles of protein conformations. FRETpredict uses an established Rotamer Library Approach to describe the FRET probes covalently bound to the protein. The software efficiently operates on large conformational ensembles such as those generated by molecular dynamics simulations to facilitate the validation or refinement of molecular models and the interpretation of experimental data. We demonstrate the performance and accuracy of the software for different types of systems: a relatively structured peptide (polyproline 11), an intrinsically disordered protein (ACTR), and three folded proteins (HiSiaP, SBD2, and MalE). We also describe a general approach to generate new rotamer libraries for FRET probes of interest. FRETpredict is open source (GPLv3) and is available at github.com/KULL-Centre/FRETpredict and as a Python PyPI package at pypi.org/project/FRETpredict.

**Author Summary:** We present FRETpredict, an open-source software to calculate FRET observables from protein structures. Using a previously developed Rotamer Library Approach, FRETpredict helps place multiple conformations of the selected FRET probes at the labeled sites, and use these to calculate FRET efficiencies. Through several case studies, we illustrate the ability of FRETpredict to interpret experimental results and validate protein conformations. We also explain a methodology for generating new rotamer libraries of FRET probes of interest.

## Introduction

Single-molecule Förster Resonance Energy Transfer (smFRET) is a well-established technique to measure distances and dynamics between two fluorophores [1, 2]. smFRET has been broadly used to study protein and nucleic acid conformational states and dynamics [3, 4], binding events [5, 6], and intramolecular transitions [7, 8]. The high spatial (nm) and temporal (ns) resolutions enable smFRET experiments to uncover individual species in heterogeneous and dynamic biomolecular complexes, as well as transient intermediates [9–13].

In a typical smFRET experiment on proteins, two residues are labeled with a donor and an acceptor FRET probe, respectively. While the FRET probes may sometimes be fluorescent proteins, they are more commonly organic molecules optimized for spectral and photophysical properties. Each such probe consists of a fluorophore and a linker, which can vary in length and is covalently attached to the protein [14]. For FRET to occur, the donor and acceptor fluorophores must have respective emission and absorption spectra that partially overlap, and the efficiency of the energy transfer depends on the proximity and relative orientation of the fluorophores.

Computational advancements, combined with enhanced sampling methods and approaches to coarse-grain, have enabled Molecular Dynamics (MD) simulations of biomolecules to explore time scales up to the millisecond or beyond [15–17].

Concomitantly, the molecular-level insights into protein structural dynamics provided by MD simulations are routinely employed to aid the interpretation of a multitude of experimental approaches, including smFRET [13, 18]. Irrespective of whether the underlying protein structure is static or dynamic, the conformational ensembles of the fluorescent probes must be taken into account to accurately predict FRET efficiencies from MD simulations [19].

To model the conformational space of dyes attached to a protein, several methods have been developed [20]. At the low end of the spectrum of computational cost, the Available Volume (AV) method uses a coarse-grained description of the probe for predicting the geometric volume encompassing the conformational ensemble of the probe [21, 22]. At the high end, MD simulations can be performed with explicit FRET probes [18, 23, 24]. Although, this approach provides unique insight into the motion of and interactions between protein and FRET probes, it must be often preceded by the parameterization of force field for the fluorescent dyes [23]. Furthermore, comparison with studies which integrate results from multiple pairs of probe positions require running independent MD simulations for each probe pair. Somewhere in the middle of the scale of computational cost and resolution is the Rotamer Library Approach (RLA), where multiple rotamer conformations of the FRET probe are placed at the labeled site of a protein conformation, and the statistical weight of each conformer is estimated using a simplified potential [25]. Polyhach *et al*. [25] introduced the RLA in the context of electron paramagnetic resonance [26]. The RLA may, however, also be employed to predict FRET [27, 28], in addition to double electron-electron resonance (DEER) and paramagnetic relaxation enhancement (PRE) nuclear magnetic resonance data [25, 26, 29, 30].

In this work we introduce FRETpredict, a Python module based on the RLA for calculating FRET efficiency based on protein conformational ensembles and MD trajectories. We describe a general methodology to generate rotamer libraries for FRET probes and present case studies for both intrinsically disordered and folded proteins (ACTR, Polyproline 11, HiSiaP, SBD2, and MalE).

## Design and Implementation

FRETpredict is written in Python and is available as a Python package. The `FRETpredict` class carries out the FRET efficiency predictions. The class is initialized with (i) a protein structure or trajectory (provided as `MDAnalysis Universe` objects [31]), (ii) the residue indices to which the fluorescent probes are attached, and (iii) the rotamer libraries for the fluorophores and linkers to be used in the calculation. The *lib/libraries.yml* file lists all the available Rotamer Libraries, along with necessary fluorophore information, including atom indices for calculating transition dipole moments and distances between fluorophores. As shown in the *Results* section, the calculations are triggered by the *run* function.

The main requirements are `Python` 3.6-3.8 and `MDAnalysis` 2.0 [31]. FRETpredict can be installed through the package manager PIP by executing

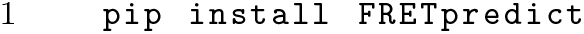

Tests reproducing FRET data for a Hsp90 system can be run locally using the test running tool `pytest`.

### Rotamer library Generation

Each FRET probe consists of two parts: the fluorescent dye, responsible for the FRET, and the linker, which comprises (i) a spacer, to distance the dye from the protein and (ii) a moiety to attach the probe covalently to the protein. For example, many of the most widely used probes can be purchased with maleimide (to link to Cys), N-hydroxysuccinimide (to link to Lys), or azide (for click chemistry) functional groups.

In FRETpredict, rotamer libraries are created through the following steps:

(i) generation of the conformational ensemble of the FRET probe using MD simulations,

(ii) selection of the peaks of the dihedral distributions of the linker, (iii) two clustering steps, and (iv) filtering. These steps are detailed in S1 Text and implemented in *FRETpredict/rotamer libraries.py*. In this work, we created rotamer libraries for AlexaFluor, ATTO, and Lumiprobe dyes with different linkers, using the force fields developed by Graen *et al*. [32]. This selection of rotamer libraries of widely used FRET probes are made available as a part of the FRETpredict package. Moreover, we provide a Jupyter Notebook tutorial (*tests/tutorials/Tutorial generate new rotamer libraries.ipynb*) which illustrates how to generate new rotamer libraries for FRETpredict.

### FRETpredict algorithm

For each protein structure to be analysed—either individually or as an ensemble—the FRETpredict algorithm places the FRET probes at the selected protein sites independently of each other. Relative orientations and distances between the dyes are then computed for all combinations of the donor and acceptor rotamers. Further, nonbonded interaction energies between each rotamer and the surrounding protein atoms are calculated within a radius of 1.0 nm. Using these energies, statistical weights are first estimated for donor and acceptor rotamers independently and subsequently combined to calculate average FRET efficiencies. S2 Text details the rotamer library placement and weighting steps. In the following, we will detail the calculation of the average FRET efficiency in different averaging regimes.

#### FRET efficiency calculation

FRET efficiency is defined as the fraction of donor excitations that result in energy transfer to the acceptor, and can be calculated as 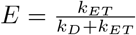, where *k_ET_* is the instantaneous FRET rate and *k_D_* is the spontaneous decay rate of donor excitation by non-FRET mechanisms (e.g. donor emission or non-radiative mechanisms). *k_ET_* can be calculated as 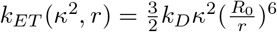, where *R*_0_ is the Förster radius, and *κ*^2^ is the orientation factor, related to the relative orientation of the dipole moments of the dyes. The Förster radius is defined as

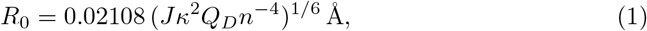

where *J* is the spectral overlap integral between the fluorescence emission of the donor and the absorption spectrum of the acceptor, *Q_D_* is the quantum yield of the donor in the absence of the acceptor, and *n* is the refractive index of the medium. Of these parameters, the most challenging to estimate is *κ*^2^. While it can be difficult to measure *κ*^2^ experimentally due to the rapid isomerization of the linker region of the probes, *κ*^2^ is often approximated to its freely diffusing isotropic average of 2/3 by considering that the fluorophore dynamics occur on a timescale that is sufficiently shorter than the donor lifetime. By assuming a fixed donor–acceptor distance, *r*, and *κ*^2^ = 2/3, we obtain

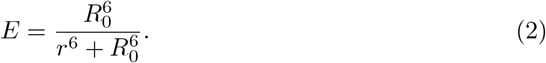

For most cases, this approximation is acceptable due to the length of the linker region and rapid fluorophore reorientation. However, the placement of the probes on a protein structure may restrict the motions of the dyes due to interactions with the surrounding protein environment. Because of such potentially restricted fluorophore motions, sometimes *κ*^2^ ≠ 2/3. Therefore, a more general formula for calculating FRET efficiency is

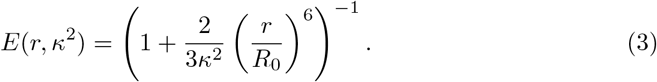

In this case, it is still assumed that the chromophore is reorienting faster than the donor lifetime, but that its motion is restricted in space. Due to the discrete nature of the RLA, FRETpredict allows precise computation of *κ*^2^ and the possibility to compute *R*_0_ in a *κ*^2^-dependent way. *κ*^2^-dependent *R*_0_ calculations (Eq. 1) are the default in FRETpredict, but users can also adopt a fixed *R*_0_ value by setting fixed_R0=True and specifying the *R*_0_ value with the r0 option. *R*_0_ values for the most common FRET probes are reported in *lib/R0/R0 pairs.csv*.

#### Averaging regimes

Protein, linker and dye motions may all contribute to FRET and so dynamics on different timescales may be important; here we simplify these as the protein correlation time (*τ_p_*), the linker-distance correlation time (*τ_l_*), the orientation correlation time of the dye (*τ_k_*), and the fluorescence lifetime (*τ_f_*). Given a conformational ensemble, but no explicit representation of the dynamical motion and timescales, the “average” FRET efficiency depends on how rapidly the various time-dependent components of *E* (i.e., *r* and *κ*^2^ in Eq. 3) are averaged relative to the fluorescence lifetime. If a specific motion occurs much faster than the fluorescence decay, the effective *k_ET_* will be completely averaged over that degree of freedom. Assuming that protein fluctuations are slow (i.e., *τ_p_ >> τ_f_*), we obtain three different regimes for the relationship between the experimentally measured efficiency and the underlying donor–acceptor distance distribution. [33]

##### Static Regime (*τ_k_ >> τ_f_* and *τ_l_ >> τ_f_*)

In this scenario, dye distance and orientation fluctuations are both slow, thus, there is no averaging of transfer rate, and every combination of protein configurations, linker distance, and dye orientation gives a separate *k_ET_*. In this case, the FRET efficiency is averaged over *N* protein conformations as well as over the *m* and *l* rotamers for the donor and the acceptor, respectively,

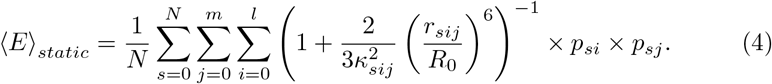

In this regime, 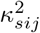 is an instantaneous value calculated for a given combination of donor and acceptor rotamers as

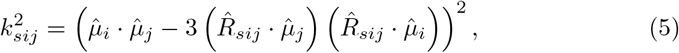

where 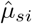 and 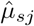 are the transition dipole moment unit vectors of the donor and acceptor, respectively, and 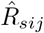 denotes the normalized inter-fluorophore displacement. In FRETpredict, the atom pairs defining 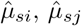, and 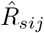 are specified in *lib/libraries.yml*.

##### Dynamic Regime (*τ_k_ << τ_f_* and *τ_l_ >> τ_f_*)

The dynamic regime is commonly assumed in the treatment of experimental data, where the complete conformational sampling is achieved within the fluorescence lifetime of the donor. The FRET efficiency is calculated as:

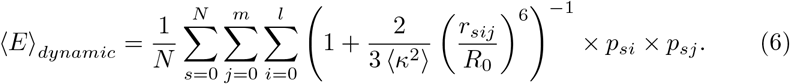

Here, 〈*κ*^2^〉 is calculated over all the protein conformations and combinations of probe rotamers:

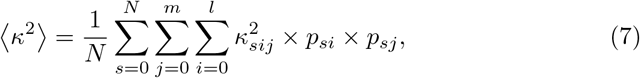

##### Dynamic+ Regime (*τ_k_ << τ_f_* and *τ_l_ << τ_f_*)

In this regime, both dye distances and orientations are very fast, and the *k_ET_* for each protein frame is averaged over all dye configurations, considering both distances and orientations. The FRET efficiency is calculated as

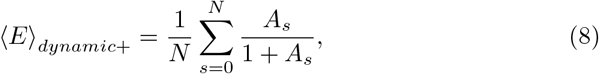

where

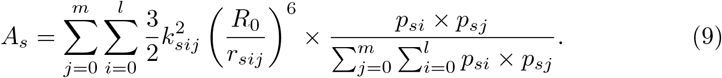

## Results

### Rotamer libraries

To illustrate the extent to which the conformational ensemble of the probes is reduced upon the generation of the rotamer libraries, we plotted the projection on the *xy*-plane of the distance vectors between the C*α* atom and the central atom of the fluorophore (Fig 2 and S6, S7, and S8 Figs) of all the generated rotamer libraries (S3, S4, and S5 Figs). Compared to the unfiltered rotamer libraries (S6 Fig), the distribution of fluorophore positions for the *large* rotamer libraries (cutoff = 10) are less isotropic and homogeneous, as evidenced by the deviation of the scatter plot from a circular shape. Unsurprisingly, the anisotropicity is increasingly more pronounced for the *medium* and *small* rotamer libraries which were obtained by filtering out cluster of less than 20 and 30 conformers, respectively (S7 and S8 Figs).

**Fig 1.**
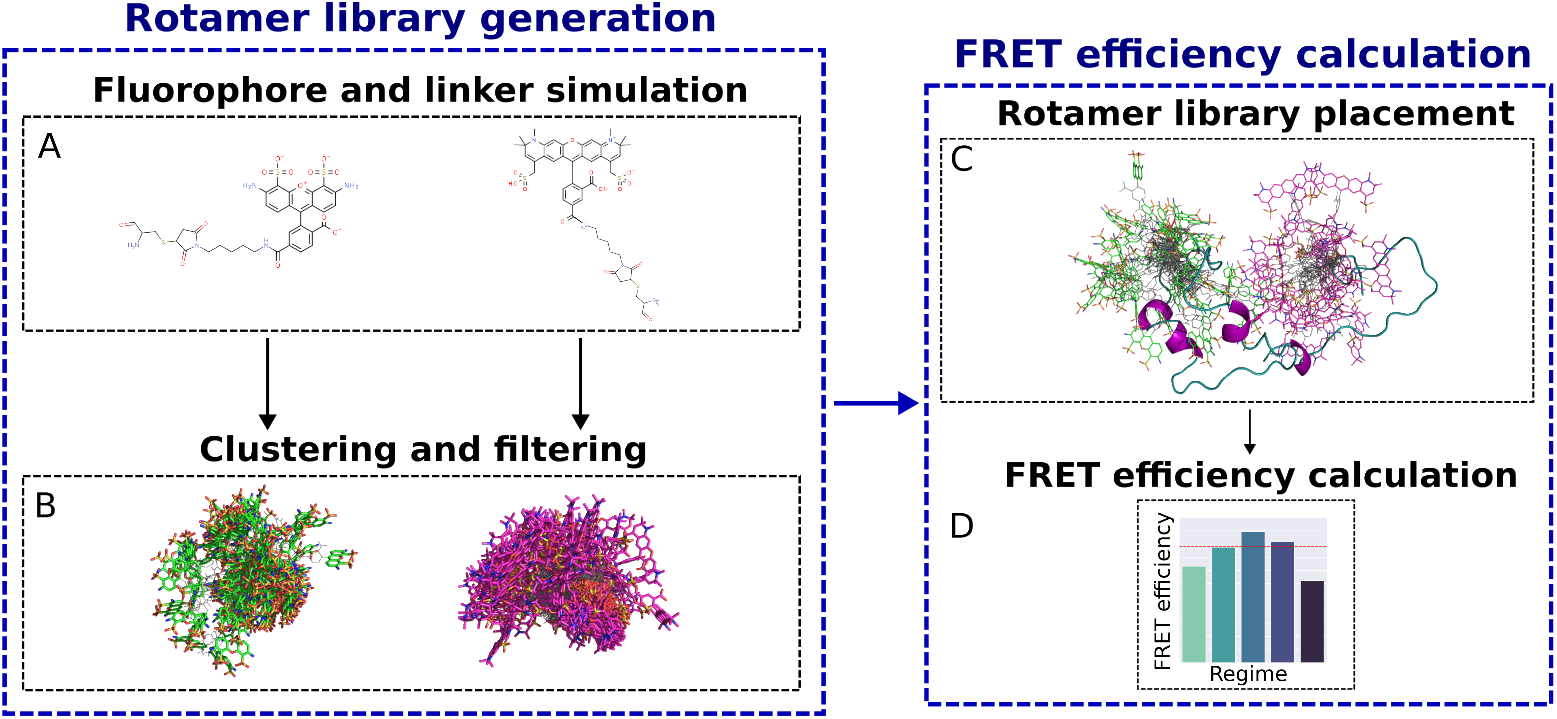
Visual summary of the functionalities in FRETpredict, which consists of two main routines: rotamer library generation (*left*) and FRET efficiency calculation (*right*). (*A*) All-atom MD simulations of a free FRET probe in solution are performed to thoroughly sample the conformational ensemble of the probe. (*B*) The obtained conformations are clustered and the clusters are filtered by population size to generate the rotamer library of the FRET probe. (*C*) The rotamer libraries of the donor and acceptor probes are placed at the labeled sites and (*D*) average FRET efficiencies are estimated according to different averaging regimes.

**Fig 2.**
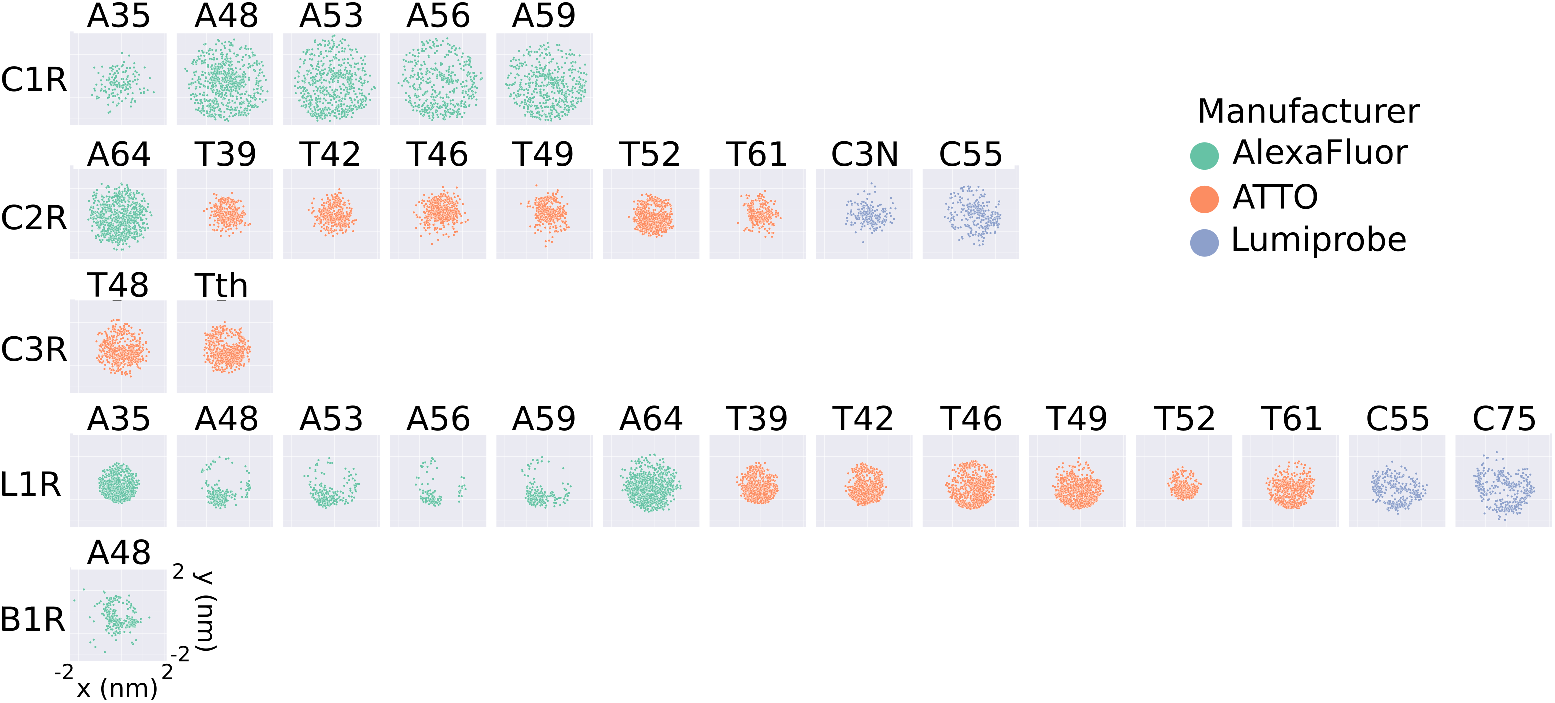
2D projections of the position of the fluorophore with respect to the C*α* atom for the *large* rotamer libraries generated in this work. The projections are obtained as the *x* and *y* coordinates of the central atom of the fluorophore (*O91* for AlexaFluor, *C7* for ATTO, and *C10* for Lumiprobe), after placing the C*α* atom at the origin. Each plot represents a different FRET probe, divided into rows according to linker type (C1R, C2R, C3R, L1R, B1R, from top to bottom), and colored according to the manufacturer (green for AlexaFluor, orange for ATTO, and blue for Lumiprobe).

The rotamer libraries of some FRET probes show pronounced anisotropy, illustrated by the deviation of the scatter plots from a circular shape (A48 L1R, A53 L1R, A56 L1R, A59 L1R, and A48 B1R). The observed anisotropy can be related to the length of the linker, and hence to its the rotational degrees of freedom. For example, the rotamer library A48 C1R is more isotropic than A48 L1R because L1R is a shorter linker than C1R (S3 Fig). On the other hand, a comparison between A48 L1R and T42 L1R suggests that the more isotropic nature of T42 might be due to the structure of the T42 fluorophore which effectively provides an extension to the linker length (S4 Fig).

The RLA relies on a trade-off between thorough conformational sampling and computational cost, as the latter increases with the increased size of the library (S9 Fig), which ideally should not exceed ~ 1, 000 rotamers. To provide an idea of the time differences involved in using rotamer libraries with different numbers of rotamers, we report the times required to calculate the FRET efficiencies for Polyproline 11 (S10 Table).

Below we showcase how FRETpredict can be used to calculate FRET efficiencies using different labels, different averaging schemes and different types and sources of protein/peptide conformations. Our goal here is not to discuss the biophysics of the individual systems, but rather to highlight the capabilities of FRETpredict.

### Case study 1: Protein Trajectory (pp11)

Polyproline 11 (pp11) has been described as behaving like a rigid rod, and was used as a “spectroscopic ruler” in the seminal paper by Stryer and Haugland [34]; subsequent work showed additional complexity [24, 33, 35, 36]. The pp11 system is thus a classical example of the importance of comparing molecular models with FRET data. Here, we compared FRET efficiency values estimated using FRETpredict with reference values from experiments [33] and from extensive all-atom MD simulations of pp11 with explicit FRET probes [23]. For analyses with the RLA we removed these FRET probes to ensure that the conformational ensembles were comparable, and thus compare the different ways of representing the dyes (explicitly or via RLA). In both experiments and simulations, the terminal residues were labeled with AlexaFluor 488 - C1R (donor) and AlexaFluor 594 - C1R (acceptor), and the *R*_0_ value was fixed to 5.4 nm. We used *large* rotamer libraries to estimate the FRET efficiency of pp11 in the three averaging regimes. Comparison with the reference values (Fig 3 and S11 Table) shows that FRETpredict calculations yield predictions that are comparable to MD simulations with explicit representation of the probes when compared with the experimental values, suggesting that the RLA provides a relatively accurate FRET calculation. In particular, the Dynamic regime best approximates the experimental value. As a convenient approach to calculate FRET efficiencies when there is no information about which averaging regime to use, we also calculate the average, ⟨*E*⟩, over the estimates of the Static, Dynamic, and Dynamic+ regimes.

**Fig 3.**
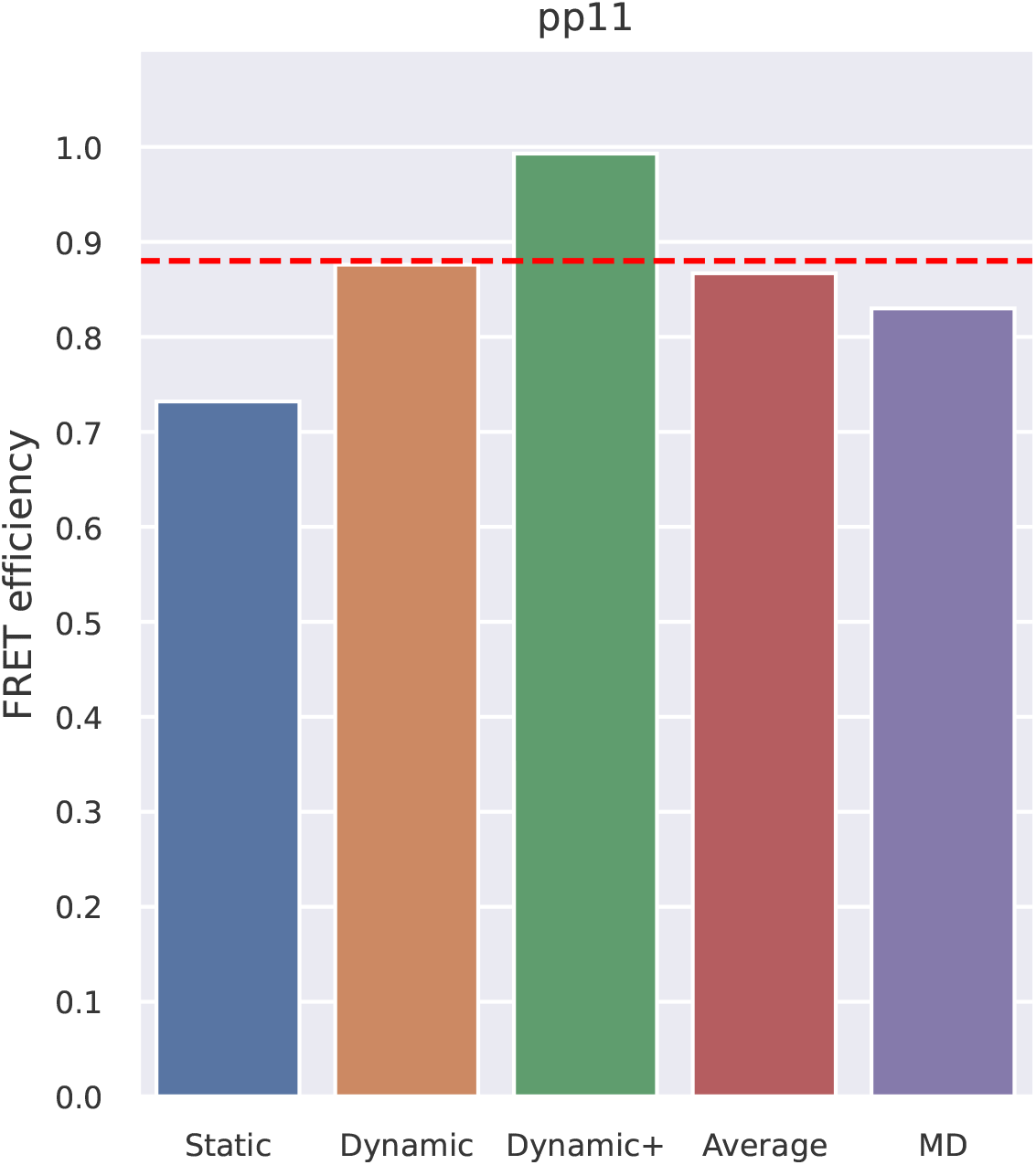
FRET efficiency obtained using FRETpredict for the MD trajectory of Polyproline 11 fluorescently labeled at the terminal residues. We calculated *E* using the *large* rotamer libraries and for the different regimes (Static, Dynamic, and Dynamic+, in blue, orange, and green, respectively). The graph also shows the average over the three regimes (Average, in red) and the *E* value obtained from MD simulation with explicit FRET probes (MD, in purple). The red dashed line indicates the experimental *E* value.

FRET efficiencies were calculated from the pp11 trajectory through the following lines of code:

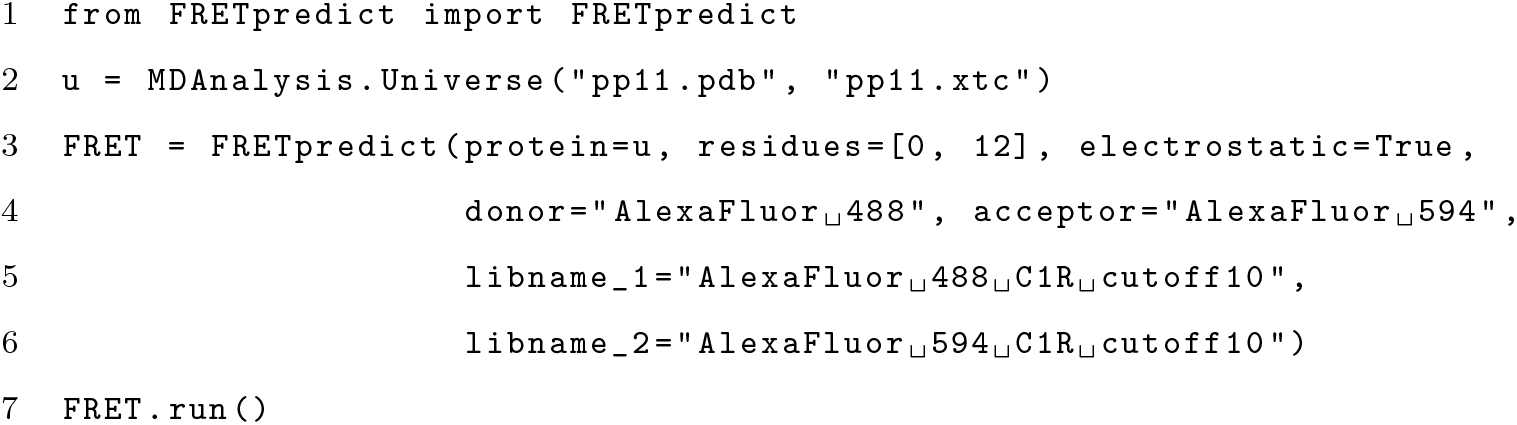

Line two generates the MDAnalysis Universe object from an XTC trajectory and a PDB topology. Line three initializes the FRETpredict object with the labeled residue numbers, the FRET probes from the available rotamer libraries, and turns the electrostatic calculations on. Line seven runs the calculations and saves per-frame and ensemble-averaged data to file. *R*_0_ was computed for each combination of FRET probes via Eq. 1.

### Case study 2: Ensemble of an Intrinsically Disordered Protein (ACTR)

ACTR (activator for thyroid hormone and retinoid receptors) is an intrinsically disordered protein that has previously been extensively studied [38, 39]. Here, we used ACTR to demonstrate how FRETpredict can be used on conformational ensembles for intrinsically disordered proteins.

We used previous experimental FRET measurements and MD simulations for ACTR solutions at different urea concentrations that were used to assess the effect of chemical denaturants on protein structure [37, 40]. As in the experiments, we labeled the residue pairs 3-61, 3-75, and 33-75 with Alexa Fluor 488 - C1R as the donor and Alexa Fluor 594 - C1R as the acceptor. To account for the dependence of *R*_0_ on urea concentration, we used Eq. 4 in Zheng *et al*. [40] and estimated *R*_0_ = 5.40 Å, 5.34 Å, and 5.29 Å for [urea] = 0 M, 2.5 M, and 5 M, respectively.

Fig 4 and S12 Table show the FRET efficiency values predicted by FRETpredict at the different urea concentrations (0 M, 2.5 M, and 5 M) using *medium* rotamer libraries. The absolute values of predicted FRET efficiency differ from the experimental values on average by 13.1, 7.2, and 12.1% for [urea] = 0 M, 2.5 M, and 5 M, respectively. Notably, the prediction trend is consistent with the experimental data for all the pairs of labeled residues of ACTR and at the three urea concentrations. The agreement between calculated and experimental trends for the *E* values shown in Fig 4 relies on the thorough and accurate sampling of conformational ensembles obtained via MD simulations by Zheng *et al*. [40] while it also contributes to validating FRETpredict as a model for calculating *E*.

**Fig 4.**
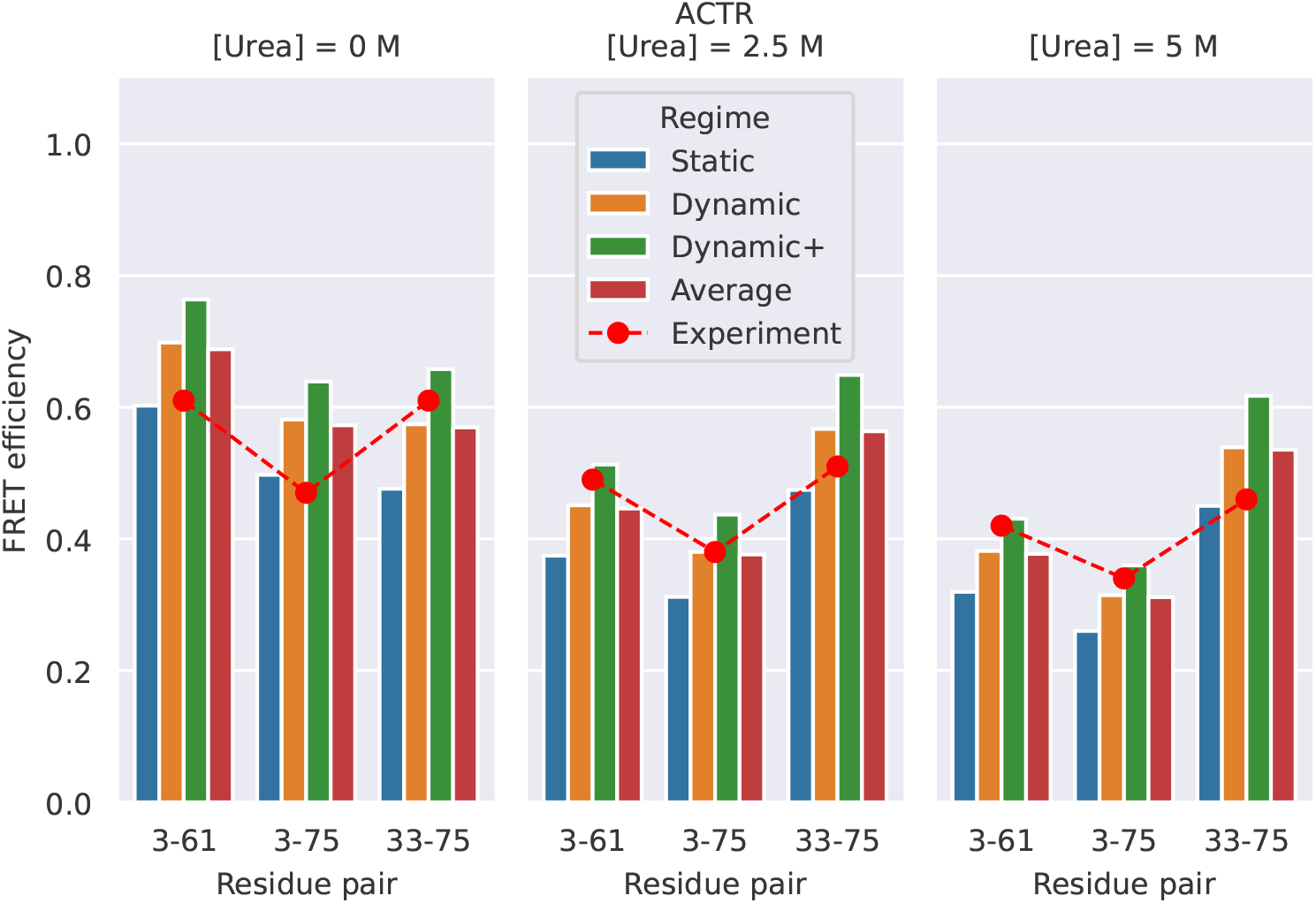
FRET efficiency for ACTR at [urea] = 0 (*left*), 2.5 M (*center*), and 5 M (*right*). The protein is fluorescently labeled at three different pairs of sites: 3-61, 3-75, and 33-75. Red circles show the experimental data from Borgia *et al*. [37]. Bars show FRETpredict estimates of the *E* values calculated using *medium* rotamer libraries. Predictions for the Static, Dynamic, and Dynamic+ regimes and their average are shown as blue, orange, green, and red bars, respectively.

To determine which regime most accurately predicts the FRET efficiency, we calculated the root-mean-square error (RMSE) between the predicted and experimental values for all the residue pairs. For the ACTR data, RMSE values obtained for the Static, Dynamic, and Dynamic+ regimes and their average are 0.233, 0.177, 0.315, and 0.171, respectively. As observed in Case Study 1, the Dynamic regime and the average best approximate the experimental FRET efficiency data.

The following lines of code were used to calculate the *E* values from the ACTR trajectory at [urea] = 0 M:

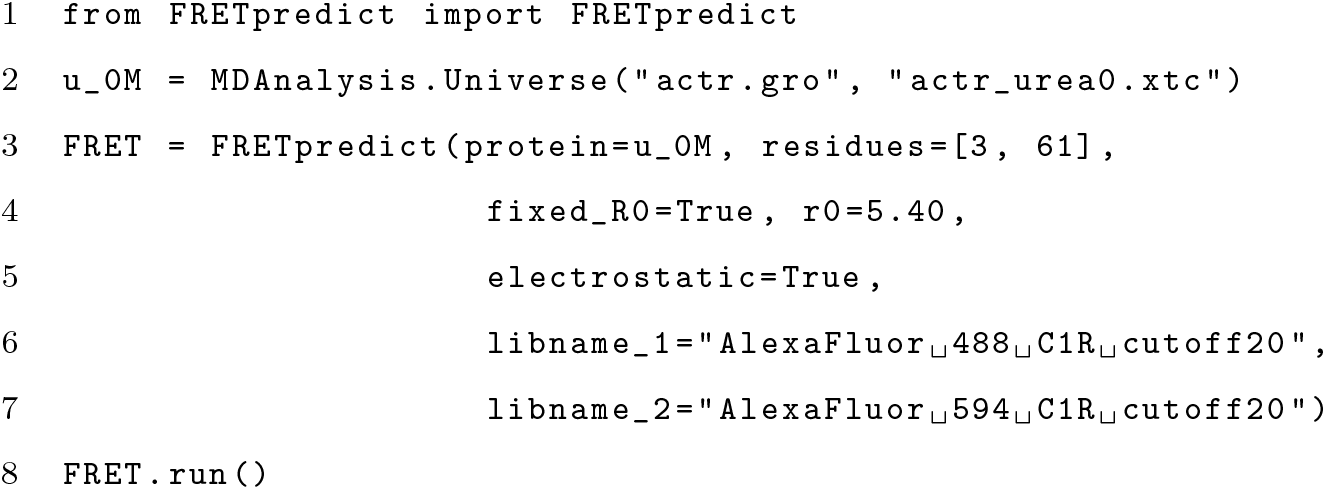

Line two generates the MDAnalysis Universe object from an XTC trajectory and a GRO topology. Line three initializes the FRETpredict object with the labeled residue numbers, the FRET probes from the available rotamer libraries, and fixes the *R*_0_ value corresponding to the specific urea concentration listed above. Line eight runs the calculations and saves per-frame and ensemble-averaged data to file.

### Case study 3: Single protein structures (HiSiaP, SBD2, MalE)

Although we generated rotamer libraries for several of the most common FRET probes, in some cases smFRET experiments might be performed with probes that are currently not available in FRETpredict. In this case study, we illustrate how, in the absence of the exact probes, accurate trends can be predicted by (i) choosing rotamer libraries with similar structural characteristics (linker length, linker dihedrals, fluorophore structure) and (ii) entering the *R*_0_ for the experimental pair of dyes (S14 Fig). We apply this strategy to the single structures of HiSiaP, SBD2, and MalE and show that it leads to results that are consistent with the experimental trends. The reference FRET efficiency data of this case study was obtained from the experimental study of Peter *et al*. [41], wherein Alexa Fluor 555 - C2R and Alexa Fluor 647 - C2R dyes were employed as donor and acceptor, respectively. In FRETpredict, both donor and acceptor were replaced by AlexaFluor 647 - C2R, the available rotamer library with the most similar steric hindrance (S3 Fig), whereas we used the *R*_0_ value of the FRET pair used in the actual experiments.

### HiSiaP

HiSiaP is the periplasmic substrate-binding protein from the sialic acid tripartite ATP-independent periplasmic transporter of *Haemophilus influenzae*. In this protein, ligand binding induces a conformational rearrangement from an open to a closed state [42].

We calculated *E* values for the labeled residue pairs measured by Peter *et al*. [41] (58-134, 55-175, 175-228, and 112-175) using structures deposited in the Protein Data Bank (PDB) for the open and closed structures (PDB codes 2CEY [43] and 3B50 [44], respectively).

The absolute values of the FRET efficiency predicted for HiSiaP differ on average by 20.6 and 24.3% from the experimental values of the open and closed conformation, respectively (Fig 5 *A* and *B*, and S13 Table). The trend of the FRETpredict prediction is about equally consistent with the experimental data for both conformations and for all the pairs of labeled residues.

**Fig 5.**
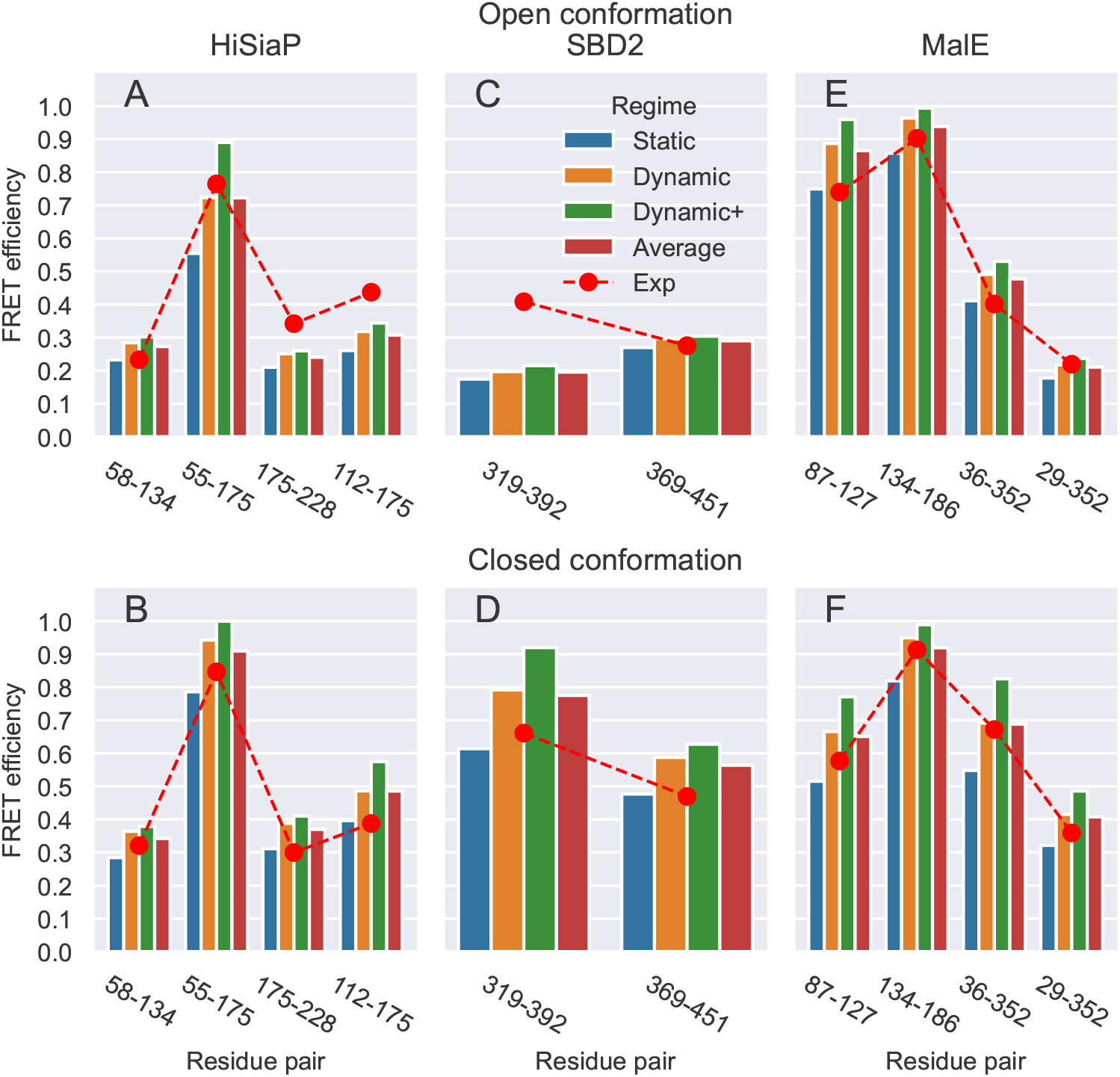
FRET efficiency values obtained on the single structures for the open and closed conformations of HiSiaP (*A* and *B*), SBD2 (*C* and *D*), and MalE (*E* and *F*) for the different residue pairs, using *large* rotamer libraries. Predictions for the Static, Dynamic, and Dynamic+ regimes and their Average are shown as blue, orange, green, and red bars, respectively. Red circles show the experimental reference values for each pair of residues.

The code used to calculate the FRET efficiency for the single HiSiaP open structure with FRETpredict is:

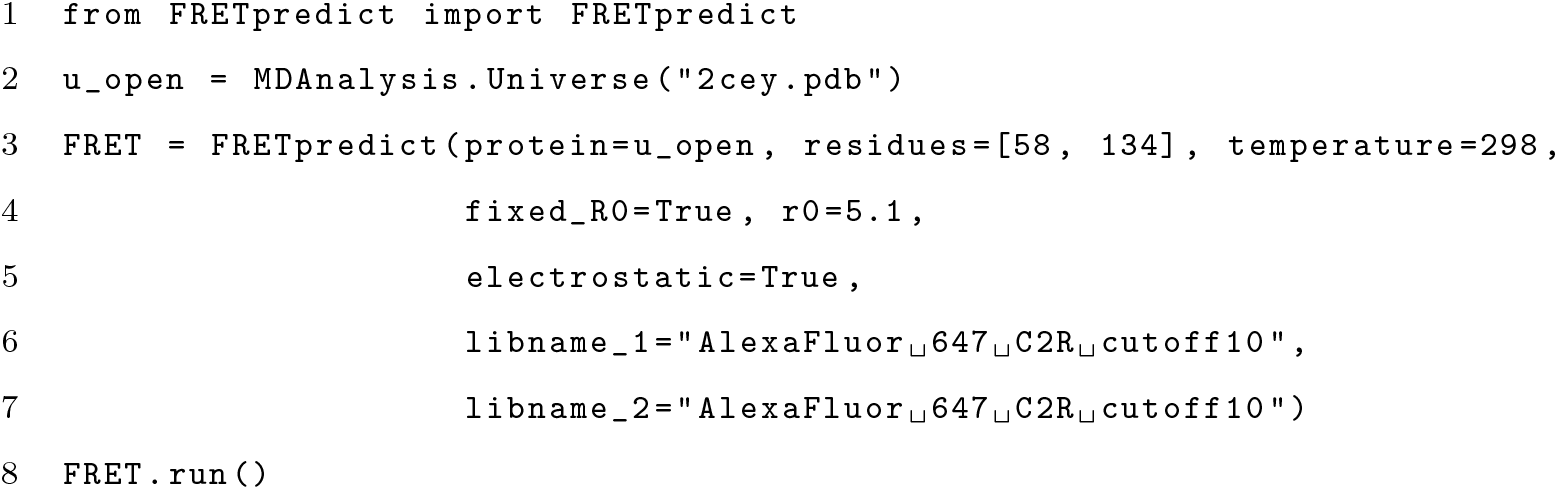

Line two generates the `MDAnalysis Universe` object for the open structure from a PDB topology. Line three initializes the FRETpredict object with the labeled residue numbers, the FRET probes from the available rotamer libraries, and fixes the *R*_0_ value to the experimental one. Line eight runs the calculations and saves per-frame and ensemble-averaged data to file. The same FRETpredict code structure has been used for the other single structure tests of SBD2 and MalE.

### SBD2

SBD2 is the second of two substrate-binding domains constituting the glutamine ABC transporter GlnPQ from *Lactococcus lactis*. As for HiSiaP, upon binding of high-affinity ligands SBD2, undergoes a transition from an open to a closed state [45].

Peter *et al*. [41] performed FRET efficiency measurements on SBD2 by labeling the residue pairs 319-392 and 369-451. We used the structures for the open and closed structures deposited in the PDB (PDB codes 4KR5 [46] and 4KQP [46], respectively) in combination with AlexaFluor 647 - C2R as both donor and acceptor.

The absolute values of the FRET efficiency predicted for SBD2 differ on average by 21.6 and 21.1% from the experimental values of the open and closed conformation, respectively (Fig 5 *C* and *D*, and S13 Table).

### MalE

The maltose binding protein from *Escherichia coli*, MalE, plays an important role in the uptake of maltose and maltodextrins by the maltose transporter complex MalFGK_2_ [47]. MalE undergoes structural transition between the apo and holo states upon sugar binding, resulting in a ~35° rigid body domain reorientation [48].

Peter *et al*. [41] performed FRET measurements on MalE by labeling the residue pairs 87-127, 134-186, 36-352, and 29-352. We used open and closed structures (PDB codes 1OMP [49] and 1ANF [50], respectively) with AlexaFluor 647 - C2R as both donor and acceptor.

The absolute values of the FRET efficiency predicted for MalE differ on average by 15.1 and 10.0% from the experimental values of the open and closed conformation, respectively (Fig 5 *E* and *F*, and S13 Table).

The RMSE values associated with the averaging regimes over all single-frame structures of HiSiaP, SBD2, and MalE are 0.097 (Static), 0.094 (Dynamic), 0.141 (Dynamic+), and 0.086 (Average). Based on these results, we observe that in the case of single-frame structures, the best predictions correspond to the Average regime.

In this case study, we used probes that are similar but not identical to those used in the experiments. The main physicochemical factors to take into consideration to assess the similarity between probes are the steric bulk of dye, the length and flexibility of the linker, and the presence of charged groups. We already noted that the steric bulk of the FRET probe and the rigidity of the linker have a strong influence on the clustering of the rotamers. Accordingly, these structural features also affect the external weights calculated upon placement of the rotamers at the binding site. On the other hand, we observed that including electrostatic interactions in FRETpredict calculations (`electrostatic=true`) had little effect on the accuracy of FRET efficiency prediction for the studied systems (S14 Fig). In summary, we found that using the rotamer library for a probe with similar steric hindrance, in combination with the *R*_0_ value for the correct dye pair, yields FRET efficiency trends in good agreement with the experimental data (S14 Fig).

## Conclusions

We have introduced FRETpredict, an open-source software program with a fast implementation of the RLA for the calculation of FRET efficiency data, along with the rotamer libraries of many of the most commonly used FRET probes. Using three case studies, we have highlighted the capabilities of our implementation in the case of a peptide trajectory (pp11), an IDP trajectory (ACTR), and single protein structures (HiSiaP, SBD2, and MalE). The FRET efficiency prediction trends are in most cases in good agreement with the experimental data; However, we note that the accuracy of the method depends on the quality and relevance of the protein conformational ensembles that are used as input.

In FRETpredict, the average FRET efficiency can be calculated in three different regimes: Static, Dynamic, and Dynamic+. We suggest using the Dynamic regime when making predictions on protein trajectories and the Static regime for single protein structures. In the absence of information about the different timescales, we find that simply averaging the results from the three regimes often leads to good agreement with experiments.

FRETpredict calculations and, more generally, FRET efficiency predictions from protein trajectories involve a trade-off between computation time and prediction accuracy. Accordingly, the choice of the optimal rotamer library selection must take its size into consideration. Large rotamer libraries may lead to greater accuracy but are also more computationally expensive than smaller libraries. On the other hand, both medium and small rotamer libraries are a good compromise between calculation time and accuracy when long simulation trajectories are used. However, using a small number of rotamer clusters may compromise the prediction of FRET efficiency, especially in case of tight placement at the labeled site, in which many rotamers may be excluded from the calculation due to probe-protein steric clashes. Therefore, we recommend using *large* rotamer libraries when the computational cost is not a limiting factor and *medium* libraries for larger conformational ensembles.

## Availability and Future Directions

The software is available on GitHub at github.com/KULL-Centre/FRETpredict, where it is published and distributed under GPL license, version 3. Tutorials for predicting FRET efficiency with FRETpredict and creating new rotamer libraries were also created and made available on the GitHub repository. FRETpredict is also distributed as a PyPI package (pypi.org/project/FRETpredict). FRETpredict has a general framework and can be readily extended to encompass non-protein biomolecules and additional rotamer libraries of FRET probes. In the current implementation, we consider all combinations of rotamers from the respective donor and acceptor libraries and independently weigh each dye based on protein-dye interaction energies, which are evaluated for the two rotamers independently. The approach could be further developed to randomly sample pairs of rotamers and to account for dye-dye interactions in the calculation of the statistical weights assigned to each pair. Further, the calculation of average FRET efficiencies could be based on the diffusive motion of the FRET probes in a potential of mean force derived from donor–acceptor distance distributions, as recently described [51] and implemented in the MMM software-tool [27].

## Acknowledgments

M.B.A.K. acknowledges funding from the Lundbeck Foundation (lundbeckfonden.com). K.L.-L. acknowledges funding via a Sapere Aude Starting Grant from the Danish Council for Independent Research (Natur og Univers, Det Frie ForskningsrÅd, 12-126214, https://dff.dk/) and the Lundbeck Foundation BRAINSTRUC initiative in structural biology (R155-2015-2666, lundbeckfonden.com). We acknowledge the use of resources at the core facility for biocomputing at the Department of Biology.

## Supporting Information

### SI: Supporting Information for “FRETpredict: A Python Package for FRET efficiency predictions using Rotamer Libraries”

#### S1 Text: Detailed description of the steps used to create new rotamer libraries

1. **Generation of the conformational ensemble of the FRET probe**. We generated conformational ensembles of the FRET probes by performing replica exchange MD (REMD) simulations, using the force fields developed by Graen *et al*. [32] with some minor corrections [52]. From these trajectories, we here saved and analysed approximately 28,000 frames.
2. **Selection of the peaks of the distributions of dihedral angles in the linkers**. We calculated the distributions of the dihedral angles in the linker using the conformational ensembles frem REMD as input. Combinations of the dihedral angles corresponding to peaks in the dihedral distributions were combined to generate distinct probe conformers corresponding to C1 cluster centers.
3. **First clustering step**. Trajectory frames are assigned to the C1 cluster centers of least-squares deviation of the dihedral angles.
4. **Second clustering step**. Averages over the dihedral angles in the trajectory frames assigned to each cluster center are calculated to generate a new set of C2 center centers. As the C2 cluster centers do not necessarily represent physical conformations of the probe, they cannot not be directly used to build the rotamer library. Instead, the probe conformation with the minimum least-squares deviation from the C2 cluster center is chosen as the representative conformation of each center. Moreover, each C2 cluster center is assigned a weight equal to the number of conformations in the cluster (cluster population). When normalized over all clusters, this statistical weight approximates the Boltzmann probability of the representative conformation for a free dye in solution, 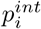. These steps are sufficient for short linkers with few dihedral angles. However, for the longer linkers in many FRET probes, extra steps are needed to decrease the number of rotamers while ensuring a good coverage of the conformational space.
5. **Filtering based on cluster populations**. In most cases, including all the C2 cluster centers into the rotamer library (e.g., 8776 conformers for Lumiprobe Cy7.5 L1R) would defeat the purpose of using the RLA as its computational cost would be considerable, albeit much lower than for an MD simulation with explicit probes. Therefore, we implemented a weight-based cutoff to reduce the number of conformations in the library while maintaining a balanced coverage of the conformational space sampled by the probes. Namely, we filtered out C2 clusters with fewer than 10, 20, or 30 members, thus obtaining new sets of C3 clusters, which will be referred in this work as *large*, *medium*, and *small* rotamer libraries, respectively. Since filtering by the assigned weights skews the remaining weights from the underlying Boltzmann distribution, we implemented a third clustering step, in which the conformations previously belonging to a discarded C2 cluster are moved to the C3 cluster of minimum least-squares deviation, and the 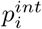 values are updated accordingly.
6. **Alignment and writing data to file**. The C3 cluster centers are aligned to the plane defined by the C*α* atom and the C–N peptide bond. The resulting rotamer library is composed of a structure file (PDB format) and a trajectory file (DCD format) for the aligned FRET probe rotamers, and a text file containing the intrinsic Boltzmann weights of each rotamer state 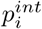.

#### S2 Text Detailed description of the rotamer library placement and weighting steps

##### Rotamer library placement

The first step in calculating FRET efficiencies is to place the FRET probes from the rotamer library at the protein site to be labeled, following the same procedure introduced in DEER-PREdict [29]. Briefly, the fluorophore library coordinates are translated and rotated based on the positions of the backbone C*α*, amide N, and carbonyl C atoms. This results in a perfect overlap with the N and C*α* coordinates of the protein backbone and an approximate alignment with the carbonyl C, which ensures that the C*α*–C*β* vector of the probe has the correct orientation relative to the side chain of the labeled residue.

##### Rotamer library weighting

For each protein conformation, the overall probability of the *i*th rotamer of a probe is estimated by combining the intrinsic and the external Boltzmann probabilities of the inserted probe, independently from the other probe. The intrinsic probabilities, 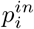, are obtained from the clustering procedure performed on the representative dihedral conformations of the free dye in solution and are related to the free energy of the rotamer, 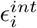, via Boltzmann inversion. Following the approach of Polyhach *et al*. [25], we account for the environment surrounding the FRET probe and calculate the probe-protein interaction energy, 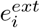. This is achieved by summing up 12-6 Lennard-Jones pair-wise interaction energies between the heavy atoms of the probe and the surrounding protein within a 1-nm radius. The Lennard-Jones atomic radii (*σ*) and potential-well depth (*ϵ*) parameters are obtained from the CHARMM36m force field [53]. The *σ* parameters can be scaled by a “forgive” factor which is set through the input parameter `sigma_scaling` and defaults to 0.5. This scaling compensates for inaccuracies in the placement of the bulky FRET probe, which tend to lead to clashes even for conformers with reasonably correct orientations of the probe with respect to the side chain of the labeled residue. The contribution of electrostatic interactions between charged probe and protein atoms is also taken into account using a dielectric constant of 78, and can be turned off by setting the `electrostatic` input parameter to `False`. Hence, the overall probability of the *i*th rotamer state attached to the *s*th protein conformation is calculated as

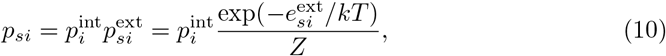

where 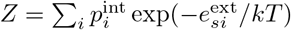 is the steric partition function quantifying the fit of the rotamer in the embedding protein conformation. Low *Z* values result from large probe-protein interaction energies, suggesting tight placement of the probe due to either (i) misplacement of the rotamers or (ii) protein conformations incompatible with the presence of the FRET probe at the labeled site. Therefore, frames with *Z* < 0.05 are discarded in the FRET efficiency calculation to preclude spurious conformers from contributing to the ensemble average, corresponding to a situation in which all of the rotamers have a positive steric energy. In FRETpredict, the default *Z* cutoff can be conveniently replaced by a user-provided value. This procedure could, in principle, be generalized to account for the effect of the probe on the protein free energy by weighting the protein conformations by the chromophore free energies −*k*_B_*T* ln(*Z*) in subsequent analysis, since the effect will differ by conformation even for those with *Z* above the cut-off.

---

**S3: Figure.**
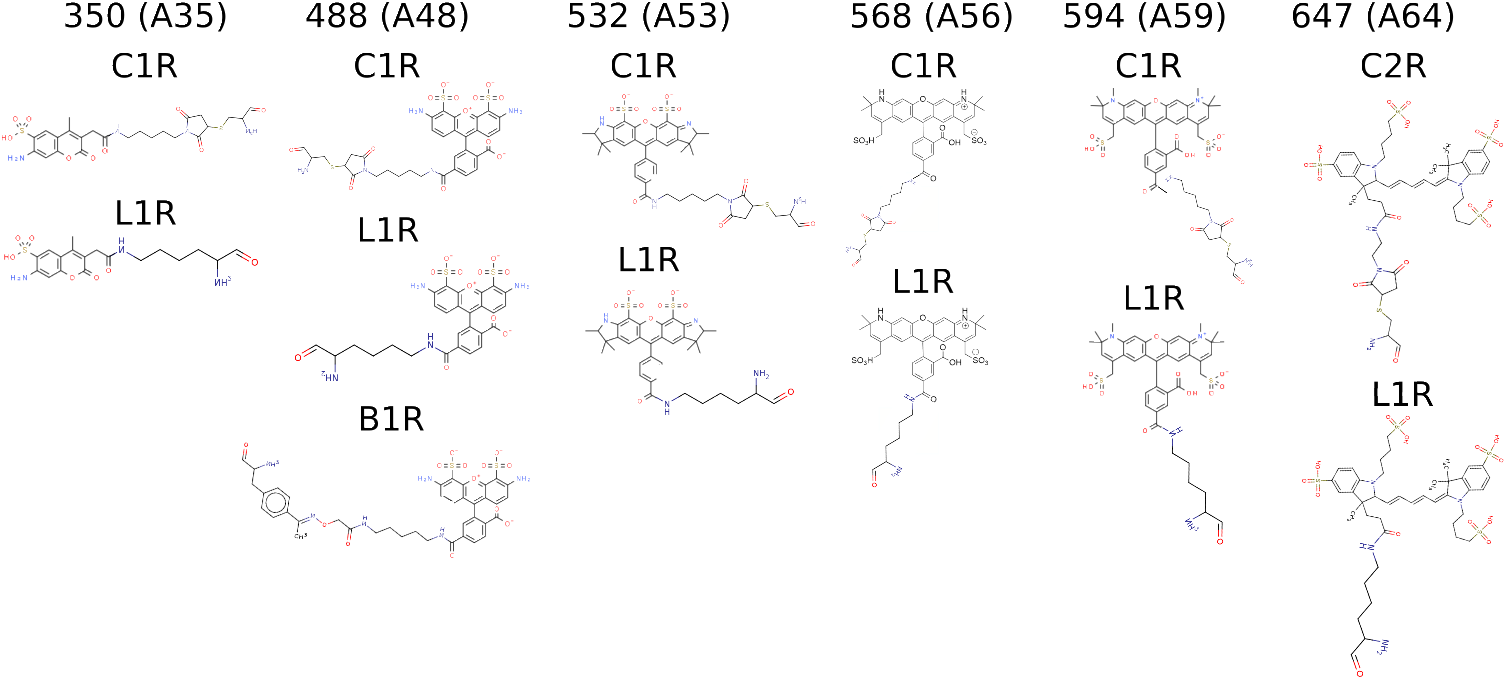
Structural formulae of the AlexaFluor probes. Structural formulae of the 13 AlexaFluor probes for which we generated rotamer libraries. Each column corresponds to a different fluorophore (acronym in parentheses). The names of the linkers are reported above each formula.

**S4: Figure.**
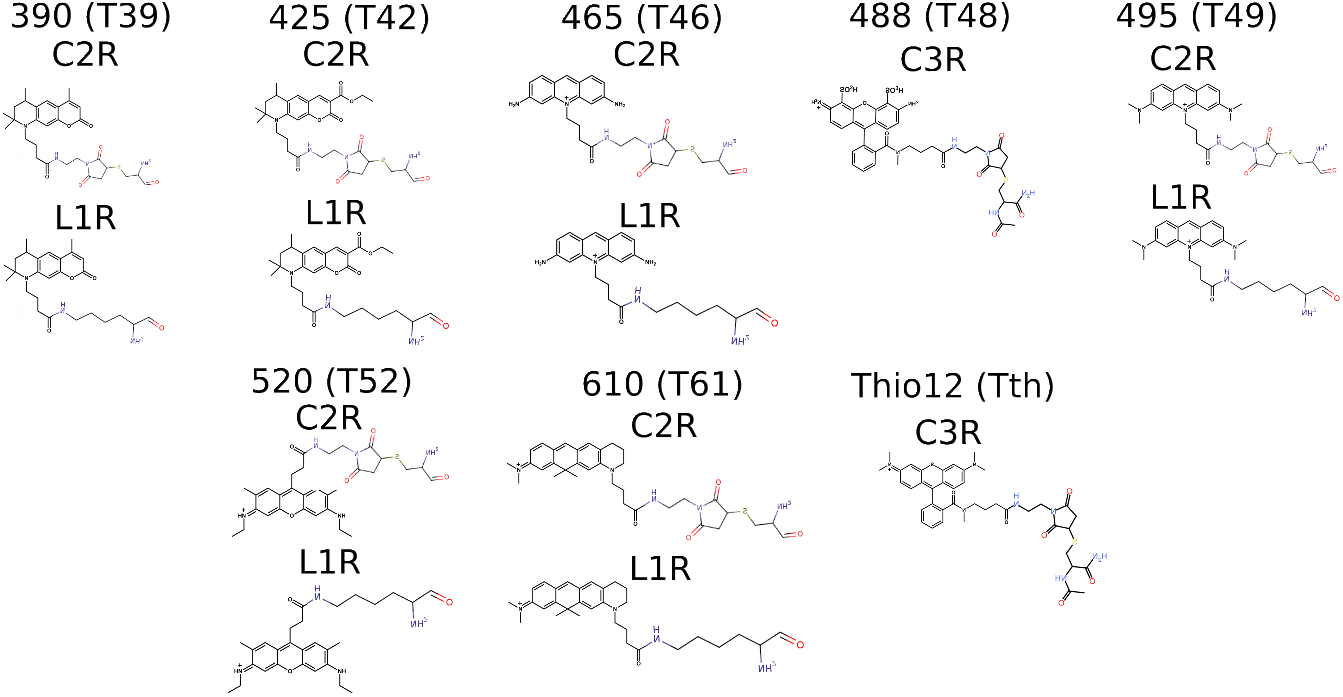
Structural formulae of the ATTO probes. Structural formulae of the 14 ATTO probes for which we generated rotamer libraries. Each column corresponds to a different fluorophore (acronym in parentheses). The names of the linkers are reported above each formula.

**S5: Figure.**
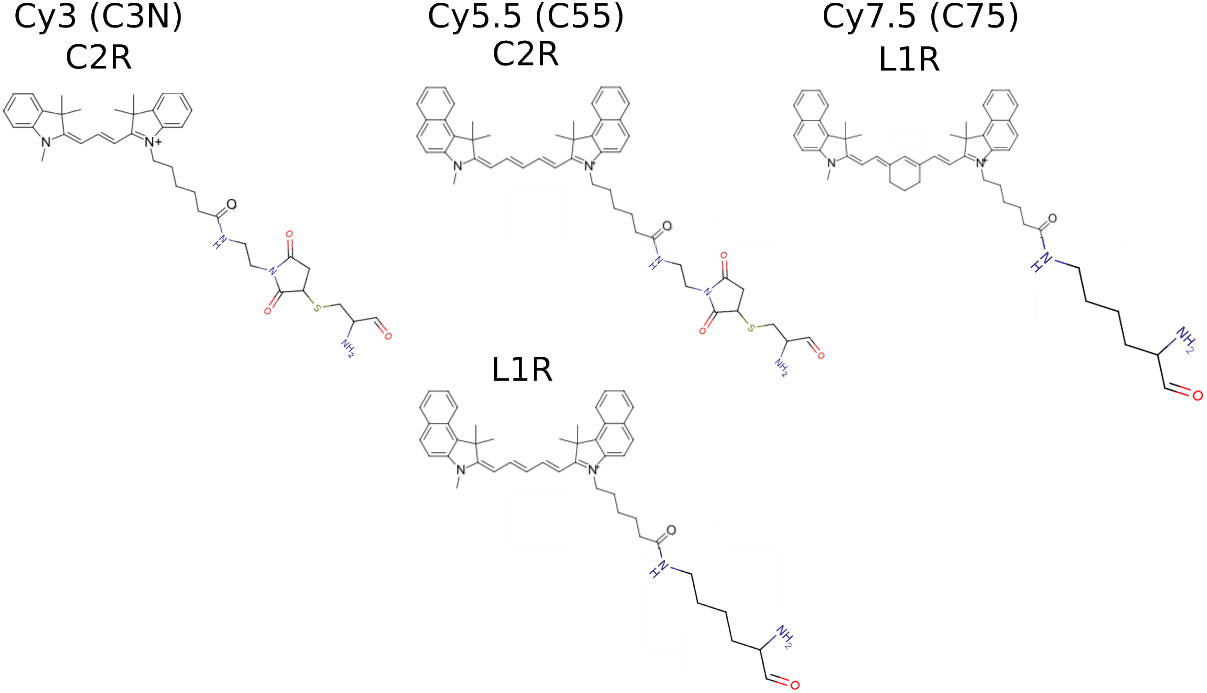
Structural formulae of the Lumiprobe probes. Structural formulae of the four Lumiprobe probes for which we generated rotamer libraries. Each column corresponds to a different fluorophore (acronym in parentheses). The names of the linkers are reported above each formula.

**S6: Figure.**
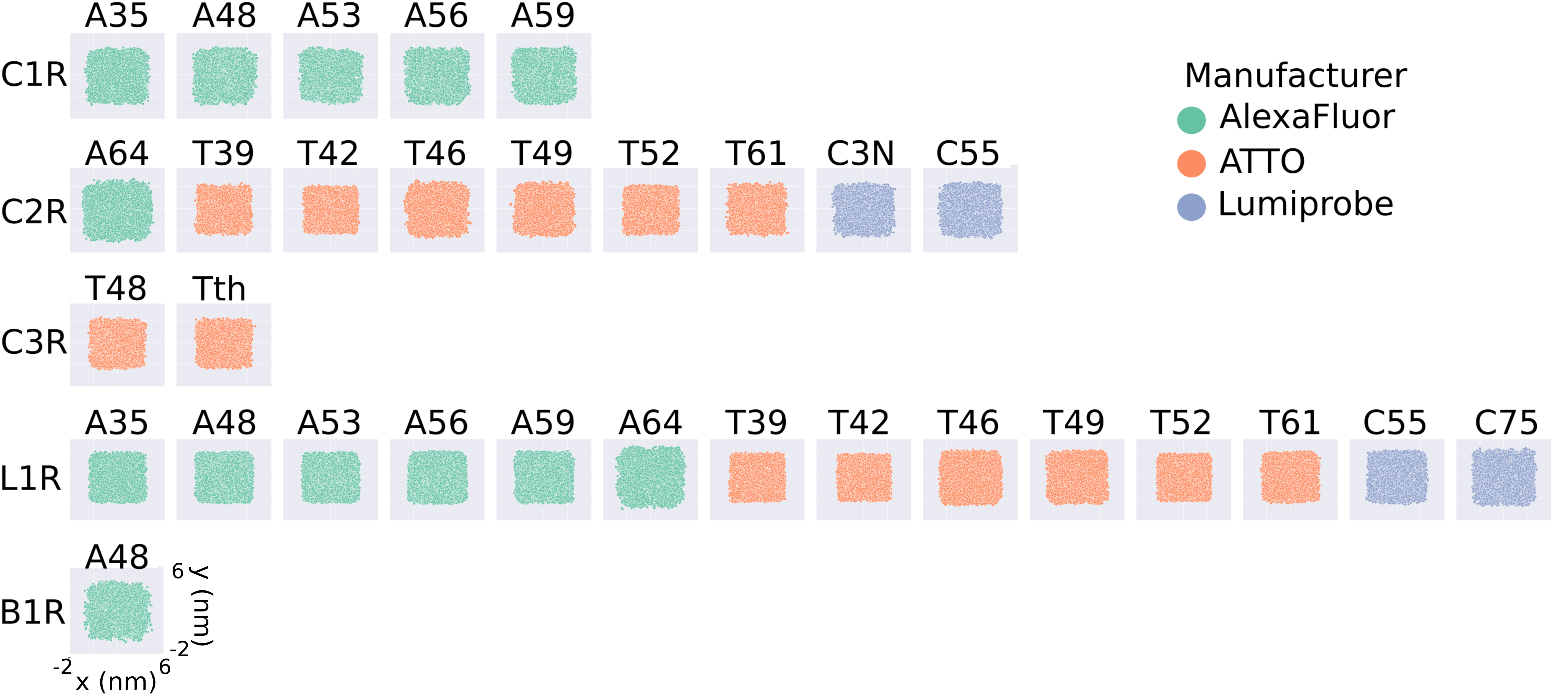
Scatter plot of Rotamer Libraries central atoms for unfiltered rotamer libraries (cutoff = 0). 2D projections of the position of the fluorophore with respect to the C*α* atom for the unfiltered rotamer libraries generated in this work (C2 cluster centers). The projections are obtained as the *x* and *y* coordinates of the central atom of the fluorophore (*O91* for AlexaFluor, *C7* for ATTO, and *C10* for Lumiprobe), after placing the C*α* atom at the origin. Each plot represents a different FRET probe, divided into rows according to linker type (C1R, C2R, C3R, L1R, B1R, from top to bottom), and colored according to the manufacturer (green for AlexaFluor, orange for ATTO, and blue for Lumiprobe).

**S7: Figure.**
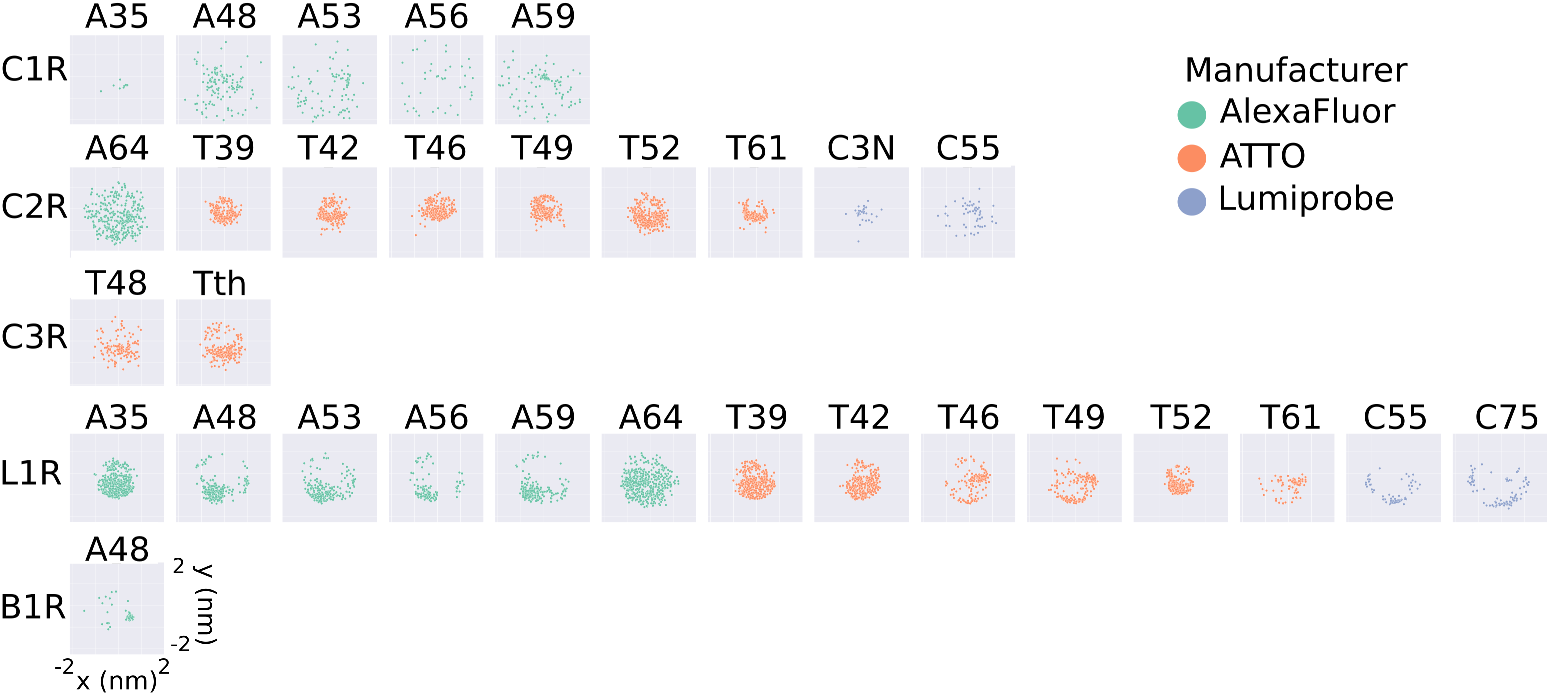
Scatter plot of medium-size rotamer libraries’ central atoms (cutoff = 20). 2D projections of the position of the fluorophore with respect to the C*α* atom for the *medium* rotamer libraries generated in this work. The projections are obtained as the *x* and *y* coordinates of the central atom of the fluorophore (*O91* for AlexaFluor, *C7* for ATTO, and *C10* for Lumiprobe), after placing the C*α* atom at the origin. Each plot represents a different FRET probe, divided into rows according to linker type (C1R, C2R, C3R, L1R, B1R, from top to bottom), and colored according to the manufacturer (green for AlexaFluor, orange for ATTO, and blue for Lumiprobe).

**S8: Figure.**
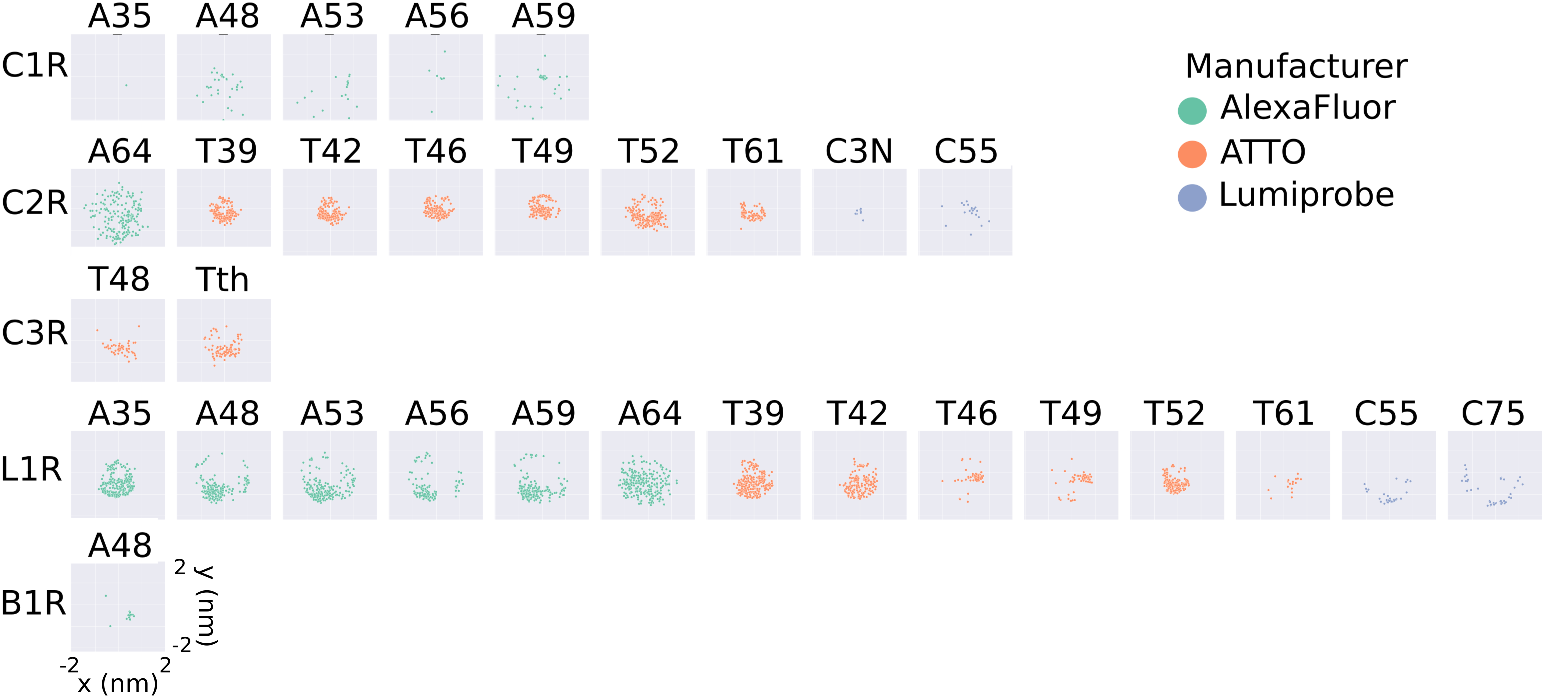
Scatter plot of small-size rotamer libraries central atoms (cutoff = 30). 2D projections of the position of the fluorophore with respect to the C*α* atom for the *small* rotamer libraries generated in this work. The projections are obtained as the *x* and *y* coordinates of the central atom of the fluorophore (*O91* for AlexaFluor, *C7* for ATTO, and *C10* for Lumiprobe), after placing the C*α* atom at the origin. Each plot represents a different FRET probe, divided into rows according to linker type (C1R, C2R, C3R, L1R, B1R, from top to bottom), and colored according to the manufacturer (green for AlexaFluor, orange for ATTO, and blue for Lumiprobe).

**S9: Figure.**
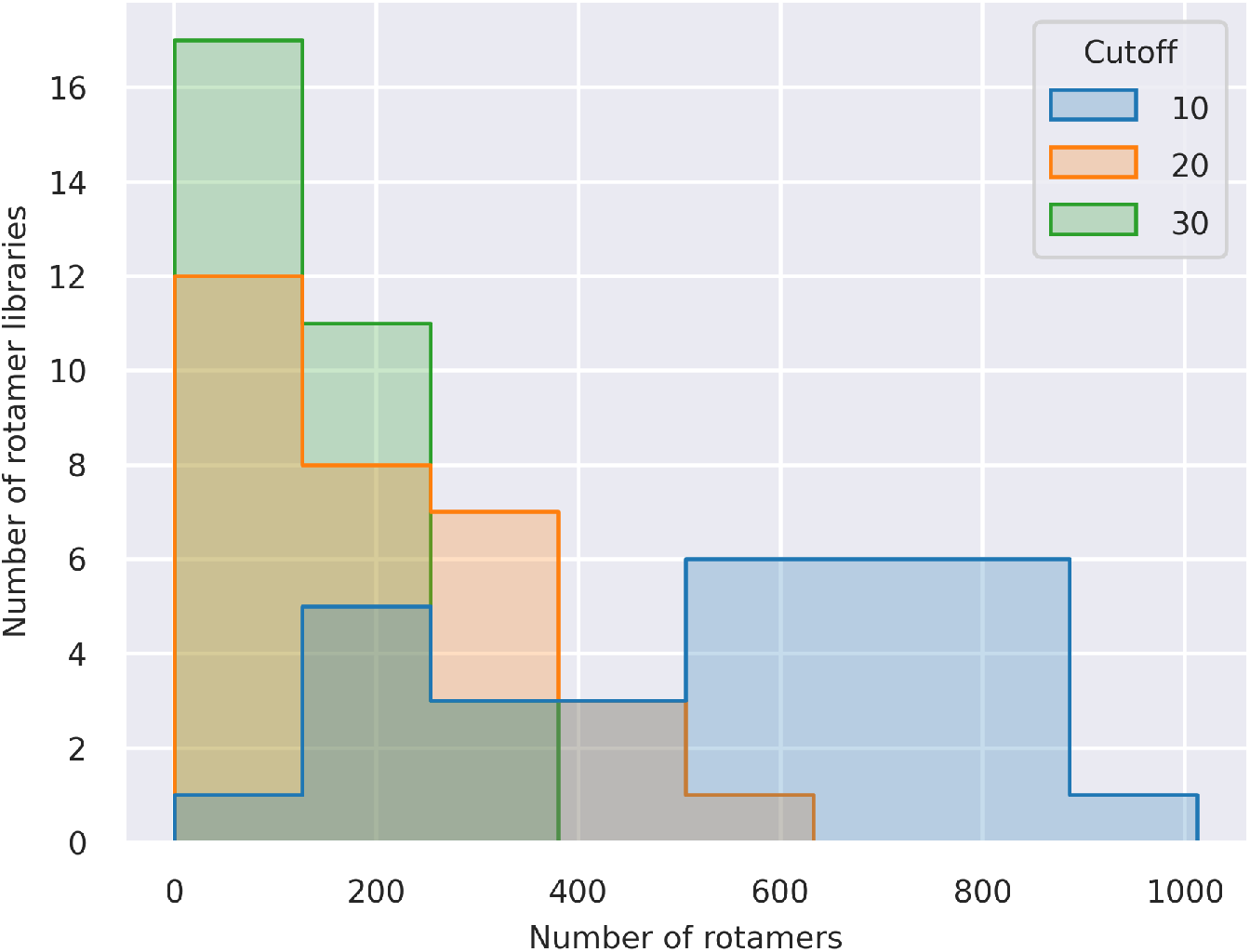
Large, medium, and small rotamer libraries populations. Distribution of the number of conformers across all the *large* (blue), *medium* (orange), and *small* (green) rotamer libraries generated in this work.

**S10 Table:** Computational times obtained using different cutoffs. Computational times required to calculate FRET efficiency from a pp11 trajectory of 316 frames (Case study 1) using the *large* (cutoffs = 10), *medium* (cutoff = 20), and *small* rotamer libraries for AlexaFluor 488 - C1R and AlexaFluor 594 - C1R, on a laptop with AMD Ryzen 7 4800h processor with a Radeon graphics card. Compared to the *large* library, the *medium* library has significantly fewer cluster centers and it lowers the computational cost by a factor 6. Instead, choosing the *small* over the *medium* rotamer library results in a gain in computation time of around a factor of 3.

|  | *small* | *medium* | *large* |
| --- | --- | --- | --- |
| Donor clusters | 706 | 124 | 32 |
| Acceptor clusters | 574 | 106 | 38 |
| Computation time | 692 s | 120 s | 37 s |

**S11 Table:** FRETpredict *E* for Case study 1: pp11 (Fig 3). FRET efficiencies calculated for pp11 using FRETpredict with different rotamer library sizes and three averaging regimes (Static, Dynamic, Dynamic+) as well as the average over those. The reference experimental value is 0.88 whereas the value obtained as the average over the three regimes from MD simulations with explicit FRET probes is 0.83.

| Polyproline 11 (pp11) |  |  |  |
| --- | --- | --- | --- |
| Regime | *small* | *medium* | *large* |
| Static | 0.732 | 0.745 | 0.743 |
| Dynamic | 0.876 | 0.886 | 0.881 |
| Dynamic+ | 0.993 | 0.972 | 0.853 |
| Average | 0.917 | 0.912 | 0.89 |

**S12 Table:** FRET efficiencies for Case study 2: ACTR (Fig 4). ACTR FRET efficiencies calculated with FRETpredict for all residue pairs (3-61, 3-75, 33-75) at different urea concentrations (0 M, 2.5 M, and 5 M), for the *medium* rotamer library. The averaging regime is reported in parentheses.

| ACTR |  |  |  |
| --- | --- | --- | --- |
| Residue pair (Regime) | [Urea] = 0 M | [Urea] = 2.5 M | [Urea] = 5 M |
| 3-61 (Exp) | 0.610 | 0.490 | 0.420 |
| 3-61 (Static) | 0.602 | 0.374 | 0.319 |
| 3-61 (Dynamic) | 0.698 | 0.451 | 0.382 |
| 3-61 (Dynamic+) | 0.763 | 0.513 | 0.431 |
| 3-61 (Average) | 0.688 | 0.446 | 0.377 |
| 3-75 (Exp) | 0.470 | 0.380 | 0.340 |
| 3-75 (Static) | 0.497 | 0.312 | 0.260 |
| 3-75 (Dynamic) | 0.581 | 0.380 | 0.314 |
| 3-75 (Dynamic+) | 0.639 | 0.437 | 0.360 |
| 3-75 (Average) | 0.572 | 0.376 | 0.311 |
| 33-75 (Exp) | 0.610 | 0.510 | 0.460 |
| 33-75 (Static) | 0.476 | 0.474 | 0.450 |
| 33-75 (Dynamic) | 0.574 | 0.567 | 0.539 |
| 33-75 (Dynamic+) | 0.658 | 0.649 | 0.617 |
| 33-75 (Average) | 0.570 | 0.563 | 0.535 |

**S13 Table:** FRET efficiencies for Case study 3: Single structure proteins (Fig 5) FRET efficiencies calculated with FRETpredict for the open and closed conformations of all the single-structure proteins (HiSiaP, SBD2, and MalE), for the *large* rotamer library. Every row corresponds to a labeled residue pair, with the protein conformation reported in parentheses. Every column corresponds to an averaging regime or to the experimental value for the specific residue pair and protein conformation.

| HiSiaP |  |  |  |  |  |
| --- | --- | --- | --- | --- | --- |
| Residue pair (conformation) | Exp | Static | Dynamic | Dynamic+ | Average |
| 58-134 (open) | 0.233 | 0.232 | 0.283 | 0.302 | 0.272 |
| 58-134 (closed) | 0.321 | 0.284 | 0.364 | 0.379 | 0.342 |
| 55-175 (open) | 0.765 | 0.554 | 0.723 | 0.890 | 0.722 |
| 55-175 (closed) | 0.847 | 0.786 | 0.942 | 0.999 | 0.909 |
| 175-228 (open) | 0.342 | 0.210 | 0.251 | 0.260 | 0.240 |
| 175-228 (closed) | 0.300 | 0.311 | 0.388 | 0.410 | 0.370 |
| 112-175 (open) | 0.437 | 0.260 | 0.318 | 0.343 | 0.307 |
| 112-175 (closed) | 0.388 | 0.396 | 0.486 | 0.575 | 0.486 |
| SBD2 |  |  |  |  |  |
| Residue pair (conformation) | Exp | Static | Dynamic | Dynamic+ | Average |
| 319-392 (open) | 0.408 | 0.174 | 0.197 | 0.215 | 0.195 |
| 319-392 (closed) | 0.661 | 0.614 | 0.792 | 0.920 | 0.775 |
| 369-451 (open) | 0.275 | 0.270 | 0.296 | 0.304 | 0.290 |
| 369-451 (closed) | 0.469 | 0.477 | 0.587 | 0.627 | 0.564 |
| MalE |  |  |  |  |  |
| Residue pair (conformation) | Exp | Static | Dynamic | Dynamic+ | Average |
| 87-127 (open) | 0.740 | 0.749 | 0.887 | 0.959 | 0.865 |
| 87-127 (closed) | 0.577 | 0.515 | 0.666 | 0.771 | 0.651 |
| 134-186 (open) | 0.903 | 0.857 | 0.964 | 0.994 | 0.938 |
| 134-186 (closed) | 0.913 | 0.819 | 0.949 | 0.989 | 0.919 |
| 36-352 (open) | 0.401 | 0.411 | 0.491 | 0.530 | 0.477 |
| 36-352 (closed) | 0.672 | 0.548 | 0.692 | 0.825 | 0.688 |
| 29-352 (open) | 0.219 | 0.177 | 0.217 | 0.237 | 0.210 |
| 29-352 (closed) | 0.359 | 0.321 | 0.415 | 0.486 | 0.407 |

**S14: Figure.**
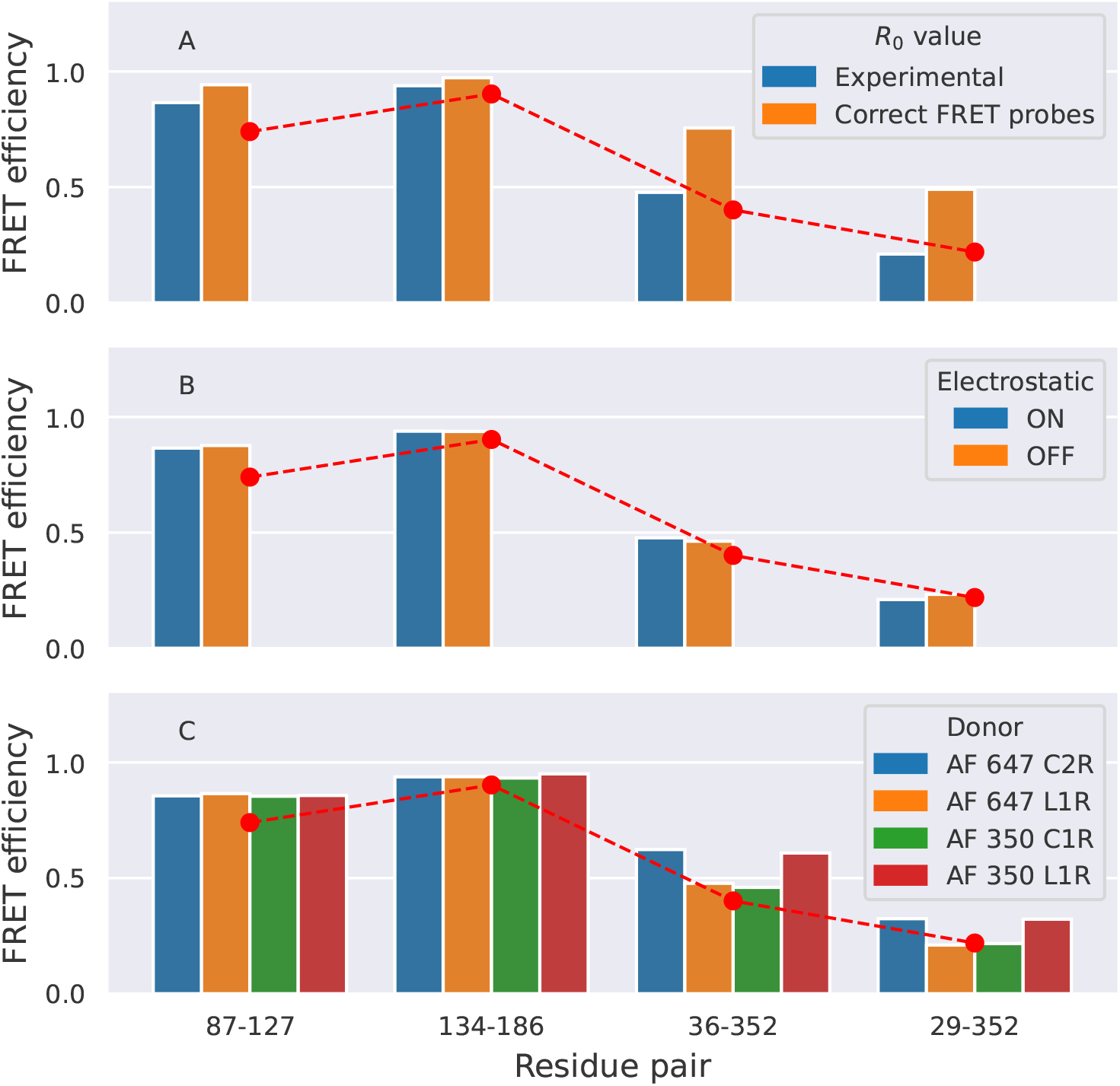
Physicochemical parameters affecting FRETpredict calculations. Effects of different physicochemical parameters on FRETpredict calculation (*R*_0_, probe steric bulk, and electrostatics, in panels A, B, and C, respectively). Calculations were performed on the open structure of MalE with the *large* rotamer library. Reported FRET efficiencies in all panels correspond to the average over the different regimes. In panel A, the *R*_0_ value is changed from the experimental value of 5.1 nm (blue bars) to the actual *R*_0_ of the two FRET probes used in the calculations (AlexaFluor 647 - AlexaFluor 647), i.e., 6.50 nm (orange bars). In panel B, the FRET efficiency was computed by turning electrostatic interactions on (blue bars) or off (orange bars) in the calculation of probe–protein energies. In panel C, the donor FRET probe is AlexaFluor 647 C2R (blue bars), AlexaFluor 647 L1R (orange bars), AlexaFluor 350 C1R (green bars), and AlexaFluor 350 L1R (red bars).

